# Disrupted modular organization of primary sensory brain areas in schizophrenics

**DOI:** 10.1101/161521

**Authors:** Cécile Bordier, Carlo Nicolini, Angelo Bifone

## Abstract

Abnormal brain resting-state functional connectivity has been consistently observed in patients affected by Schizophrenia (SCZ) using functional MRI and other neuroimaging methods. Graph theoretical methods provide a framework to investigate these defective functional interactions and their effects on the modular organization of brain connectivity networks. A few studies have shown abnormal distribution of connectivity within and between functional modules, an indication of imbalanced functional segregation ad integration in SCZ patients. However, no major alterations in the modular structure of functional connectivity networks in patients have been reported, and unambiguous identification of the neural substrates involved remains elusive. Recently, it has been demonstrated that current modularity analysis methods suffer from a fundamental and severe resolution limit, as they fail to detect features that are smaller than a scale determined by the size of the entire connectivity network. This resolution limit is likely to have hampered the ability to resolve differences between patients and controls in previous cross-sectional studies. Here, we apply a novel, resolution limit-free approach to study the modular organization of resting state functional connectivity networks in a large cohort of SCZ patients, and in matched healthy controls. Leveraging these important methodological advances, we find new evidence of substantial fragmentation and reorganization involving primary sensory, auditory and visual areas in SCZ patients. Conversely, frontal and prefrontal areas, typically associated with higher cognitive functions, appear to be largely unaffected, with changes selectively involving language and speech processing areas. Our findings provide support to the hypothesis that cognitive dysfunction in SCZ may arise from deficits occurring already at early stages of sensory processing.

## Introduction

Schizophrenia has been associated with aberrant functional connectivity as measured by neuroimaging methods in a number of studies (Friston & Frith 1995; Liang et al. 2006; Liu et al. 2008; Calhoun et al. 2009; Karbasforoushan & Woodward 2012; Garrity et al. 2007). This growing evidence is in keeping with the disconnectivity hypothesis of Schizophrenia (Friston & Frith 1995) that posits that the core dysfunction of this disease may correspond to alterations of the functional interactions between specialized brain areas (Bullmore et al. 1998; Ellison-Wright & Bullmore 2009; Fornito et al. 2009; Kubicki et al. 2005), resulting in defective integration of activity in distributed networks and in cognitive disintegration (Tononi & Edelman 2000). Indeed, psychotic symptoms akin to those of schizophrenia, including hallucinations and delusions, are also observed in certain neurological disorders that involve disruption of corticocortical and corticosubcortical connections (Hyde et al. 1992; Su et al. 2015; Bullmore et al. 1998). Understanding the nature of connectivity alterations in SCZ patients and their effects on brain functional integration may provide important insights into the etiology of this devastating disease, as well as potential diagnostic or prognostic markers.

To this end, graph theoretical approaches have been proposed as a powerful framework to assess topological features of functional connectivity networks (Bassett & Bullmore 2006; Bullmore & Sporns 2009; Kaiser 2011; Stam & Reijneveld 2007; Reijneveld et al. 2007), in which nodes correspond to anatomically defined brain regions and the edges to interregional correlations. Several alterations in graph-related metrics of resting state connectivity have been identified in schizophrenia patients, including reduction in global network efficiency (Bassett et al. 2008; Liu et al. 2008; Bullmore & Sporns 2009), small worldness (Anderson Ariana & Cohen 2013; Yu et al. 2011; Liu et al. 2008) and rich-club organization of high-connectivity nodes (van den Heuvel et al. 2013).

Recently, graph analyses of resting state brain connectivity networks have been applied to study the brain modular organization, i.e. the presence of functionally segregated module, or "communities", within large-scale, integrated functional connectivity networks (Salvador et al. 2005; He et al. 2009; Meunier et al. 2009). Typically, these methods assess patterns of edges in the graph to identify clusters of nodes that are more densely connected, denoting stronger interactions among themselves than with the rest of the system. This mathematical formulation embodies the notion of segregation and integration, as the emergence of modules reflects the balance between intra- and inter-cluster connections. Hence, community detection methods enable investigation of the interplay between functional segregation and integration in the healthy and diseased brain, and provide a means to map the brain’s modular organization. Changes in the structure of specific modules in patients may highlight specific circuits or neural substrates affected by the disease. Moreover, modularity analyses make it possible to identify the brain connector hubs, i.e. the regions that are responsible for the integration of the activity of the modules, and to assess the effects of disease on these hubs (Heuvel & Sporns 2013; Crossley et al. 2014). Indeed, there is growing evidence that abnormalities in nodes characterized by high topological centrality and connectivity are implicated in several neuropsychiatric disorders, and that connector hubs may present increased vulnerability (Crossley et al. 2014).

Several studies have investigated the modular structure of resting state functional connectivity networks derived from functional MRI in Schizophrenia patients compared to healthy controls (Alexander-Bloch et al. 2010; Alexander-Bloch et al. 2013; van den Heuvel et al. 2010; Fornito et al. 2012; Lo et al. 2015; Lerman-Sinkoff & Barch 2016; Yu et al. 2012; Liu et al. 2008). Reduction in Modularity, a measure of segregation of functional modules within an integrated network, was found in Childhood Onset Schizophrenia (Alexander-Bloch et al. 2010). However, no strong evidence of group differences in the dispersion and structure of brain modules was found in that study (Alexander-Bloch et al. 2010). Reduced Modularity was associated with a proportional increase in inter-cluster edges and decrease in intra-cluster edges in patients (Alexander-Bloch et al. 2012). Lerman-Sinkof et al. (Lerman-Sinkoff & Barch 2016) reported similar modular structures in adult schizophrenia patients and healthy subjects under stringent control of potential sources of imaging artifacts, with small but significant alterations of node community membership in specific patients networks. Yu et al. (Yu et al. 2012) found reduced overall connectivity strength and a larger, even though very limited, number of communities in the patients’ group (6 in SCZ subjects vs 5 in healthy controls). These pioneering investigations provide important indications that the strength of division of resting-state functional connectivity networks into modules may be altered in patients affected by schizophrenia. However, the partitions *per se*, i.e. the clustering of different brain regions into modules, appear very coarse, comprising only a few, broad modules that are similarly distributed in patients and controls. Hence, it remains unclear whether schizophrenia affects specific brain functional subsystems, or is just associated with a global alteration in the balance between segregation and integration.

Graph theory as applied to the study of brain networks is still in its infancy, and several methodological and conceptual issues that are still open may have affected early studies. An important finding in complex network theory is that most community detection methods, like those applied in previous studies in Schizophrenia patients, suffer from a resolution limit (Fortunato & Barthélemy 2007), as they cannot resolve clusters of nodes that are smaller than a scale determined by the size of the entire network. This limit, first demonstrated for Newman’s Modularity, is quite general and affects, to a different extent, all methods that seek to identify the community structure of a network through the optimization of a global quality function (Newman 2006), including Reichardt and Bornholdt’s (Reichardt & Bornholdt 2006), Arenas and Gomez’ (Arenas et al. 2008), Ronhovde and Nussinov’s (Ronhovde & Nussinov 2010), Rosvall and Bergstrom’s (Rosvall & Bergstrom 2008; Kawamoto & Rosvall 2015) and others. The introduction of a resolution parameter in the quality function has been proposed as a means to improve detection of smaller clusters (Alexander-Bloch et al. 2010; Reichardt & Bornholdt 2006). However, this approach introduces a specific scale determined by the choice of parameter values (Thomas Yeo et al. 2011; Reichardt & Bornholdt 2006; Ronhovde & Nussinov 2010), enabling detection of smaller clusters at the expense of larger ones, which may be unduly subdivided, resulting in partitions with relatively uniform cluster size distributions (Lancichinetti & Fortunato 2011).

We have recently demonstrated (Nicolini & Bifone 2016; Nicolini et al. 2017) that the resolution limit severely hampers the ability to resolve the modular organization of human brain connectivity networks, and to capture their complex community structure. This pervasive limit is likely to have biased previous studies in clinical populations, and may have prevented detection of differences in the organization of functional connectivity in patients and controls at a finer scale. Indeed, even though previous studies in SCZ populations systematically report substantial changes in functional connectivity and modularity strength compared to healthy controls, differences in the number, size and boundaries of functional modules appear to be modest and inconsistent across studies, often dependent on the specific clustering approach that was adopted. The deleterious effects of the resolution limit propagate to the evaluation of important topological parameters that depend on the network’s community structure. These include the participation coefficient, a parameter that enables the identification of highly-connected nodes, or hubs, responsible for the integration and efficient exchange of information between modules (Bullmore & Sporns 2009). These limitations have made it difficult to unambiguously identify the neurofunctional substrates involved in what is sometimes regarded as a disconnectivity syndrome, and to assess different hypothesis on its etiology. Defective functional interactions may be widespread and affect overall efficiency of the network (Liu et al. 2008; Bullmore & Sporns 2009; Bassett et al. 2008), or involve more specific circuits, including fronto-hippocampal (Meyer-Lindenberg et al. 2005), fronto-parietal (Garrity et al. 2007), and thalamo-cortical connections (Woodward et al. 2012). On the other end of the spectrum, it’s been hypothesized that the complex symptomatology of SCZ may arise from local deficits within primary sensory cortices (Daniel C Javitt 2009), and that impairment of higher cognitive functions may result from a bottom-up propagation of these deficits. Overcoming the limitations of current methods would help discriminate between these different scenarios and assess the relative merits of different theories underlying the disconnectivity hypothesis in schizophrenia.

Recently, we have shown that Surprise, a fitness function rooted in probability theory (Nicolini & Bifone 2016), behaves as a resolution-limit free function for community detection. Extension of this method to weighted networks, dubbed Asymptotical Surprise, was validated in synthetic and real world networks, revealing a heterogeneous modular organization of the human brain, with a wide distribution of clusters spanning multiple scales (Nicolini et al. 2017). The improved resolution afforded by Surprise makes it possible to appreciate differences in the structures of networks from different groups that are undetectable by resolution limited methods (Nicolini & Bifone 2016), and has led to a refinement of the classification of brain hubs (Nicolini et al. 2017).

Here, capitalizing on these important methodological advances, we apply Asymptotical Surprise to resolve and compare the modular structures of resting state functional connectivity networks in two cohorts of 78 schizophrenia subjects and 91 controls. In contrast with previous studies, we find profound changes in the resting state brain connectivity structure of schizophrenia patients, with a substantial functional reorganization reflecting both fragmentation and merging of functional modules. Additionally, we investigate alterations in node-wise participation coefficients and the resulting rearrangement of brain integrative regions in patients. The ability to resolve these changes at a finer scale than previously possible sheds new light on the functional implications of aberrant functional connectivity in Schizophrenia.

## Materials and Methods

### Participants

MRI data from 78 patients with schizophrenia strict diagnosis (DSM IV) SCZ (64 males, 14 females) and 91 healthy controls (CON) (65 males, 26 females) were downloaded from the open COBRE database ({http://fcon_1000.projects.nitrc.org/indi/retro/cobre.html)(Ambite et al. 2015; Wang et al. 2016). Age ranged from 18 to 65 years in both groups.

All subjects in the COBRE were screened and excluded if they had history of neurological disorder, history of mental retardation, history of severe head trauma with more than 5 minutes loss of consciousness, history of substance abuse or dependence within the last 12 months. Ethical statements are contained in the original publication of this dataset (Çetin et al. 2014). Patients were treated with one of the three atypical antipsychotics, olanzepine, risperidone or ziprasidone, and had retrospective and prospective clinical stability. None of the patients was under mood stabilizing treatment at the time of study.

### fMRI acquisition and pre-processing

Images were acquired with a Siemens MIND TRIO 3T scanner equipped for echo-planar imaging (EPI). Echo-planar imaging was used for resting state fMRI data collection with (Repetition Time) TR=2s, (Echo Time) TE=29ms, matrix size: 64×64, slices=33, voxel size=3×3×4 mm^3^ (for more details see (Çetin et al. 2014)). A total of 150 volumes of functional images were obtained for all subjects except one (this subject was excluded from the present study).

The data were pre-processed using SPM8 (Wellcome Trust Centre for Neuroimaging, London, UK). After discarding the four initial volumes, the remaining volumes were slice-timed, head-motion realigned (translational displacement and rotation along and around X, Y and Z axes) and normalized to the standard MNI EPI template space (voxel-size re-sampled to 3×3×3 mm^3^). No significant differences were observed in head-motion realignment parameters between the two groups (see Discussion and Supplementary Information sections).

### Adjacency matrices and connectivity parameters

For each participant, 638 regional mean time series were computed by averaging the voxel time series within each of the parcellized areas of the template described in (Crossley et al. 2013). After regressing movement parameters, we estimated the connectivity matrix by computing pairwise inter-regional correlation for each individual. In order to compute group-level connectivity matrices, individual correlation matrices were Fisher-transformed and averaged by group. Several critiques of correlation as a measure of functional connectivity have been proposed (Smith et al. 2011), but this definition is the most widely accepted choice, and we have adopted it to ensure comparability of our results with previous studies.

From the adjacency matrix, we extracted the distribution of z-score (corresponding to the weighted edges of our network), and we computed nodal and global measures of connectivity. The degree of a node represents the number of connections towards the other nodes. The network density indicates the ratio between the connections in the matrix and all possible connection (Albert & Barabási 2002).

Global efficiency represents the average of the inverse shortest weighted path lengths connecting every pair of nodes, and is inversely related to the network characteristic path length (Rubinov & Sporns 2010). The weighted global efficiency can be written as:

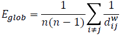

Where 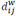 is the weighted shortest-path length between node *i* and node *j* of a graph with *n* nodes.

Weighted local efficiency quantifies a network’s resistance to failure on a local scale. The definition of weighted local efficiency used in this manuscript is the one given by (Rubinov & Sporns 2010).

### Asymptotical Surprise

The quality function that we chose to determine community structure in this study is Asymptotical Surprise (Nicolini et al. 2017), a recently developed approach rooted in information theory that aims to encode the relative entropy between the observed intracluster density and its expected value as on the basis of the Erdos-Renyi null model (Traag et al. 2015). Asymptotical Surprise was recently shown to be quasi-resolution-limit free, and to provide improved means to resolve the modular structure of complex networks of brain functional connectivity (Nicolini & Bifone 2016; Nicolini et al. 2017). Asymptotical Surprise is defined as:

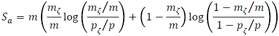

where *m* is the sum of all edge weights, *m*ζ is the sum of intracluster edge weights, *p* is the total number of possible links, *p*ζ is the total sum of possible intracluster links.

Optimization of Asymptotical Surprise was carried out by means of PACO (PArtitioning Cost Optimization), an iterative agglomerative algorithm built on a variation of the Kruskal algorithm for minimum spanning trees (Nicolini & Bifone 2016; Nicolini et al. 2017). We have recently shown that maximization of Asymptotical Surprise enables detection of heterogeneously distributed communities (Nicolini et al. 2017), thus making it possible to resolve differences in the modular organization of different networks representing functional connectivity in different subjects or experimental groups (Nicolini & Bifone 2016). A MATLAB toolbox including binary and weighted versions of Surprise optimization is available upon request at http://forms.iit.it/view.php?id=68447. PACO is a non-deterministic heuristic, like many other methods used to maximize quality functions for community detection. This means that multiple runs of PACO on the same graph may yield slightly different resulting partitions. In this study, we used 1000 runs and picked the partition with the highest value of Asymptotical Surprise. As shown in (Nicolini et al. 2017), Asymptotical Surprise does not suffer from degeneracy of nearly optimal solutions, and is characterized by a sharply distinct global optimum. Hence, there is no need to determine a consensus solution over an ensemble of different nearly optimal partitions with similarly large values of the fitness function; here, the solutions with the highest value of Asymptotical Surprise was selected.

### Percolation Analysis

In weighted networks, sparsification procedures are often applied to remove the weakest edges, which are the most affected by experimental noise, and to reduce the density of the graph, thus making it theoretically and computationally more tractable. To this end, we used a percolation analysis approach. This method, grounded in statistical physics, was first demonstrated in brain networks by Gallos et al. (Gallos et al. 2012), and previously applied to study phase transitions of connected subgraphs in random networks (Callaway et al. 2000; Goerdt 2001). In brief, percolation analysis measures the size of the largest connected component of the network upon iterative removal of the weakest edges and enables data-driven determination of the optimal sparsification threshold that preserves network structure and connectedness while removing potentially spurious correlations.

In the Erdos-Renyi random graph, the size of the largest connected component shows a sharp transition at some threshold value (Callaway et al. 2000). Contrarily, brain networks exhibit multiple percolation thresholds, revealing a hierarchy of clusters (Gallos et al. 2012). Based on these observations, we have recently demonstrated that a threshold just above fragmentation of the largest connected component strikes the optimal balance between removal of noise-induced spurious correlations and discarding of information that may be contained in the weaker links (Bordier et al. 2017)(Nicolini et al. 2017). Specifically, the percolation threshold maximizes information that can be extracted about the network’s modular organization, simultaneously maximizing sensitivity and specificity in the assignment of nodes to different modules, as shown in synthetic networks endowed with a ground-truth community structure (Bordier et al. 2017) and in human resting state functional connectivity networks (Nicolini et al. 2017). Thresholding by percolation analysis has been previously applied in human (Gallos et al. 2012) and animal (Bardella et al. 2016) functional connectivity studies.

Percolation analysis was performed independently in the two groups prior to community detection by Asymptotical Surprise maximization. This is an important conceptual step, as it overcomes a conundrum in the comparison between groups characterized by different connectivity strengths. In many previous studies, thresholds were determined by fixing the same edge densities in the connectivity graphs of the groups to be compared. However, constant edge density may bias group comparisons when the experimental groups exhibit intrinsic differences in connectivity strength, like in the present case. Imposing equal densities for graphs describing connectivity in patients and controls may lead to the inclusion of a greater number of potentially spurious links in the group with weaker connectivity, and to the exclusion of important links in the group with stronger connectivity. A higher proportion of spurious connection results in a more random network topology, and intergroup differences may just reflect different levels of noise, rather than genuine topological differences (Heuvel & Fornito 2014). Identification of optimal thresholds that maximize information about the modular organization of each group enables unbiased comparison of the two community structures. It should be noted that modules are defined in terms of node membership and do not dependent on the total density of edges, but rather on the balance between inwards and outwards edges.

### Group level comparison

After community detection by Asymptotical Surprise in the two populations (SCZ and CON), we computed the similarity between the extracted modular partitions in terms of normalized mutual information (NMI) (Danon et al. 2005), and used an approach proposed by Alexander-Bloch et al. (2012) to test for statistical differences. This method is based on the idea that if variance in the community structure data is reliably explained by group membership, then the mean NMI between all possible pairs of participants within an experimental group should be higher than the mean NMI of pairs of participants from random groups. As the underlying distribution of group mean NMI is unknown, a null-distribution is generated through a permutation method between the two experimental groups (10000 permutations).

### Participation coefficient

To complete the investigation at the node level, we considered the alteration in node role between the two populations based on the differences in modular organization revealed by Asymptotical Surprise. To this end, we adopted Guimera’ and Amaral classification scheme (Guimerà & Amaral 2005), whereby nodes are classified by their within-community degree z (a measure of how well connected a node is to other nodes in the same community) and their participation coefficient *P*, a parameter that reflects the extent to which a node is connected to nodes in other modules. *P* can be written as:

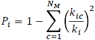

where *k*_*ic*_is the number of links of node *i* to nodes in module *c*, and *k*_*i*_ is the total degree of node *i*. To assess statistical differences, we computed the *P* index of each node for each subjects and ran a t-test between groups, Bonferroni corrected. Node-wise statistical significance was parametrically mapped on the MRI template.

## Results

### Weaker connectivity in SCZ patients

To examine the difference in functional connectivity between the two populations, and to relate this study to previous ones, we compared group-averaged pairwise correlations. Fig.1A. shows the edge-weight distributions for the two groups. We observed a significant shift in the average z-score value of the matrix between the patient and the control. Fig.1B. shows the distribution of node degree, a node-wise parameter that measures the number of edges incident to each node. A significant decrease of average node-degree was observed for the SCZ groups (green line) compared to the CON group (black line), with degree values of 25.73±13.04 and 53.83±40.35 respectively. Additionally, we computed the global efficiency coefficient *E*_*glob*_, related to the inverse average path length connecting any two nodes, a measure of how efficiently information is exchanged across the network. *E*_*glob*_ was strongly reduced in the SCZ group, with a value of 0.23 versus 0.32 for the control group, a further indication of reduced network integration in the patient.

**Figure 1.**
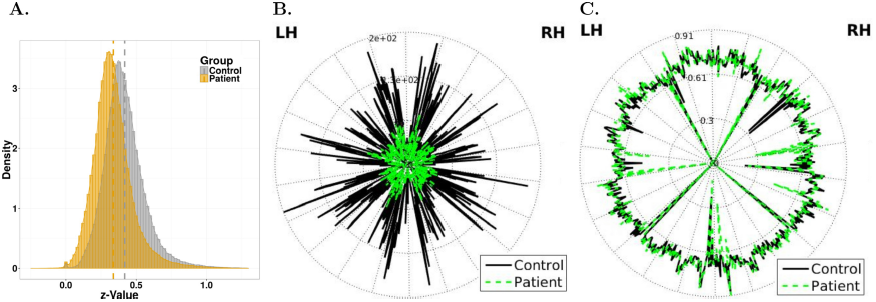
A.Z-value distribution of the adjacency matrices for the two experimental groups; a left shift in the distribution for the SCZ group indicates overall weaker connectivity. B. Degree of each node for the patients’ and control groups (in green and black, respectively). C. Local efficiency value by nodes (same color scheme as in B). The nodes of the left and on the right hemisphere (LH and RH) are respectively on the right and on the left side of the circle.

Altogether, these analyses show widespread alterations in functional connectivity in the Schizophrenia group, entirely consistent with previous reports (Liu et al. 2008; Alexander-bloch et al. 2010; Alexander-Bloch et al. 2013; van den Heuvel et al. 2010; Fornito et al. 2012; Lo et al. 2015; Lerman-Sinkoff & Barch 2016; Yu et al. 2012). Interestingly, though, local efficiency, defined as the efficiency of a node’s local network of nearest neighbors when the node is removed, was similar or even higher in the schizophrenia group 0.6±0.09 in controls; 0.68±0.09 in patients). Fig.1C. shows local efficiency for each node, with green and black lines denoting patients and controls, respectively (t-test *p-value= 0.103*). The graph displays only minor differences between the two populations, thus suggesting that network tolerance to node faults is generally preserved in patients compared to controls even if overall functional connectivity is substantially weaker (Lynall et al. 2010). This has been observed in previous studies, and pointed to as a potential evolutionary advantage that may justify persistence of schizophrenia-related genes in the general population (Lynall et al. 2010).

In summary, all graph-related parameters measured in this study appear consistent with those reported in the literature for smaller groups or specific sub-populations of patients (e.g. childhood onset schizophrenics), thus corroborating the idea that the present data-set has global and node-wise functional connectivity features that are comparable to those of previous studies. In the following, we investigate the effects of these scale-dependent differences on the modular organization of functional connectivity in schizophrenia patients vs controls using our novel community detection approach.

### Fragmentation of primary sensory areas, but not prefrontal regions, in SCZ patients

In order to determine the optimal modular partitions for our experimental groups, we applied maximization of Asymptotical Surprise by PACO, a resolution-limit free method that we have recently demonstrated in healthy volunteers (Nicolini et al. 2017). Prior to community detection, the group level adjacency matrices were sparsified using a percolation analysis approach to remove weaker edges and reduce the effects of noise, thus maximizing information about the network’s modular structure (Alexander-Bloch et al. 2010; Nicolini et al. 2017; Gallos et al. 2012; Bardella et al. 2016). Fig.2 shows the group-level adjacency matrices, with the node indexes rearranged by module membership, for the control and schizophrenia groups. Disjoint clusters of nodes, or modules, are delineated by red lines. We found 44 communities in the control group, with module sizes ranging between 141 and 1 nodes.

**Figure 2.**
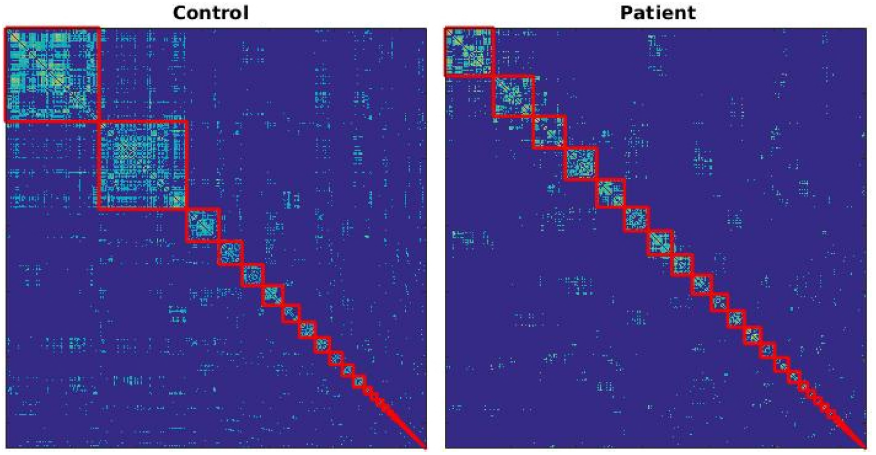
Region-by-region correlation matrices sorted by module membership obtained by Asymptotical Surprise for the two populations. Modules are demarcated by a red line. The number of modules is 44 and 39 for the control and patient group, respectively.

The optimal partition of the patients’ group comprised 39 communities, and showed a less heterogeneous size distribution (ranging between 73 and 1 nodes). The statistical significance of the difference in community structure was assessed using a recently proposed permutation approach based on Normalized Mutual Information (Alexander-Bloch et al. 2012), resulting in a *p-value= 0.009* (1000 permutations, false discovery rate corrected). The smaller number of modules in the schizophrenia group may appear somewhat counterintuitive, in the light of overall weaker functional connectivity strength in this group. However, while some of the larger modules appear to break up into smaller modules in patients, the tail of the distribution of community sizes is fatter in the schizophrenia group, thus indicating aggregation and reorganization of smaller modules. The overlaps between modules in the optimal partitions for the two groups are shown and discussed in the Supplementary Information section (Fig.S1).

Fig.3 shows the distribution of functional modules in the two groups overlaid on an anatomical template. Note that the colors denoting the communities were chosen independently in the two groups to maximize contrast between adjacent modules. Differences in the modular structures of functional connectivity in the two groups are apparent, and involve complex reorganization of nodal membership across modules. To facilitate visual inspection and interpretation of differences between the two groups, we show some communities of the control group individually, and the overlapping communities in the patients’group (Fig. 4, 5and 6).

**Figure 3.**
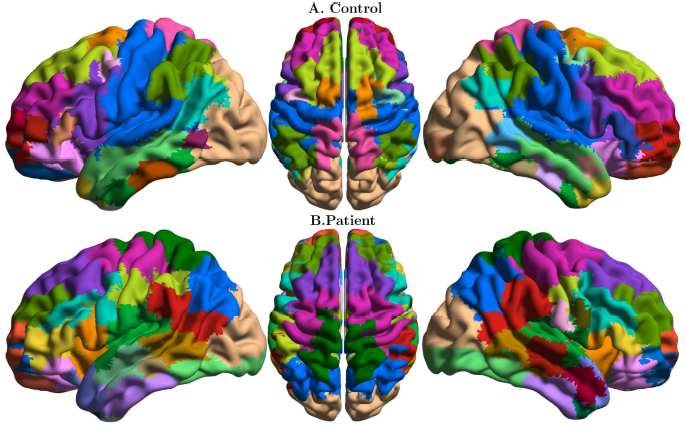
Functional connectivity modules obtained by Asymptotical Surprise overlaid on an anatomical template for: A. healthy controls; B. schizophrenia patients; colors were assigned independently in the two groups to maximize contrast between adjacent modules.

**Figure 4.**
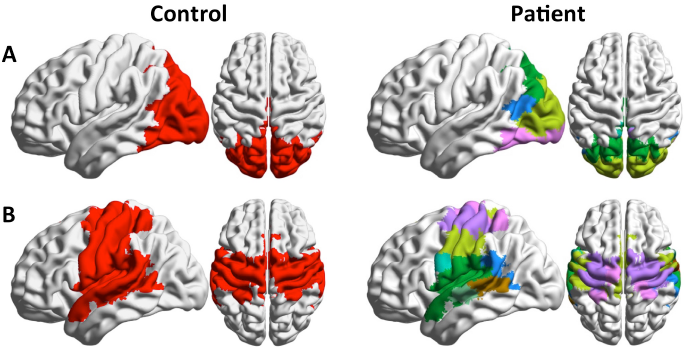
Community 1 and 2 (A and B, corresponding to visual and sensorimotor modules, respectively) of the control group (left), and overlapping communities in the patient group (right). These modules are substantially fragmented and reorganized in the SCZ group compared to healthy controls.

**Figure 6.**
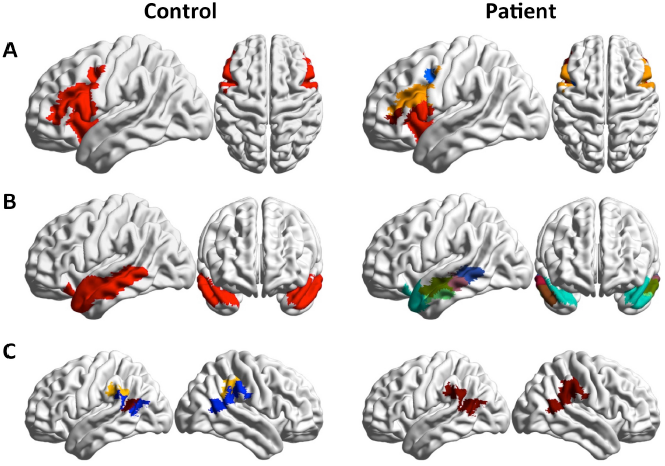
A: Community 4 of the control group and overlapping communities in the patient group (right). The Broca Area breaks apart in patients and forms an independent community with increased centrality (see also Fig. 6). B: Control’s community 7, corresponding to the medial temporal gyrus, is fragmented rostrocaudally in SCZ patients. C: the supramarginal and angular gyrus represent separate communities in controls, and merge into the same module (community 14) in the patient’s group.

The main differences in modular organization between the two groups involve the sensorimotor, visual and auditory cortices. The controls’ large occipital module (Community 1, Fig. 4A) is split in the patients’ group, with primary visual cortex standing as an independent community together with the caudal part of the inferior temporal gyrus. The more dorsal part of the occipital community includes a portion of the superior parietal lobule in healthy controls, but not in SCZ patients, where the boundary of this community lies in the vicinity of the parieto-occipital fissure. The large central module in the healthy controls’ group (Community 2, Fig.4B) comprises somatosensory, sensorimotor cortices and temporal auditory cortices, consistent with previous findings in healthy volunteers (Nicolini et al. 2017). In schizophrenia patients, this module breaks up dorsoventrally into four different clusters of nodes.

Conversely, the local modular structure of prefrontal areas is consistent between the two groups, with similar size and anatomical distribution of node membership (Fig. 5). However, long-distance connectivity between prefrontal and parietal regions is reduced in SCZ patients (Fig.5A), resulting in a separation of the Lateral Parietal Cortices from the Default Mode Network. Between-group differences in frontal lobe organization pertain particularly the language regions, with the Broca area forming an independent community in patients (Fig. 6A).

**Figure 5.**
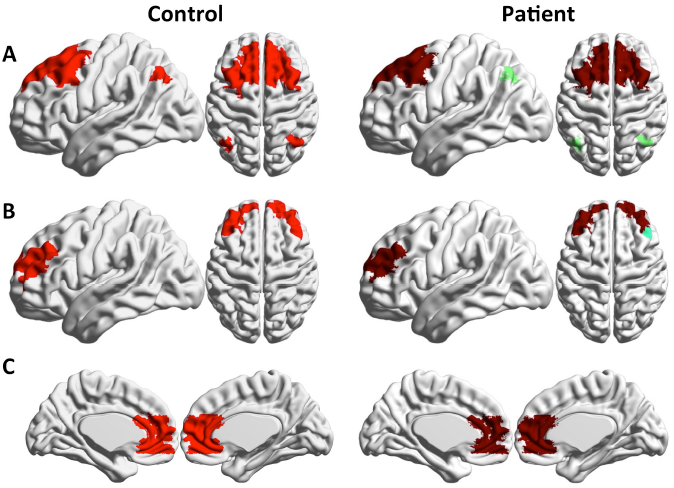
Community 3, 6 and 11 (A,B and C, corresponding to prefrontal and frontal modules) of the control group (left), and overlapping communities in the patient group (right). Local modular structure appears consistent between the two groups, with reduction in long-range fronto-parietal connectivity is observed in Community 3.

Substantial fragmentation and reorganization is also observed in the modular structure of the temporal cortex. In controls, we find well-delineated modules comprising the middle temporal gyrus and the inferior temporal gyrus, while the superior temporal gyrus is part of larger community that includes somatomotosensory cortices. In patients, the superior temporal gyrus is separated from the larger somatosensory community, and is split into two modules, anterior and posterior, respectively. The middle temporal gyrus (Community 7 Fig.6B) is split rostrocaudally into 4 different communities that include parts of the superior and inferior gyri. The inferior gyrus consists of two modules. The anterior module includes part of the middle temporal gyrus, while the posterior one extends to the primary visual cortex. Interestingly, the angular gyrus and the supramarginal gyrus appear as separate modules in healthy controls, but in patients these areas are merged into a single community including the temporoparietal junction (see Fig.6C).

In summary, Asymptotical Surprise reveals substantial alterations of the modular structure of functional connectivity in SCZ patients, with fragmentation of sensory and sensorimotor areas, and reorganization of language related areas. Conversely, prefrontal areas show similar local organization in patients and controls.

### Significantly increased centrality of sensory areas in SCZ patients

Fig.7 shows statistically significant differences in participation coefficient, a measure of diversity in intermodular connections of individual nodes. Nodes characterized by high participation coefficients have many links pointing to different modules, and are thought to play an integrative role. Nodes with most links pointing to other nodes within the same community are dubbed provincial hubs, and contribute to defining functional segregation of their communities.

**Figure 7.**
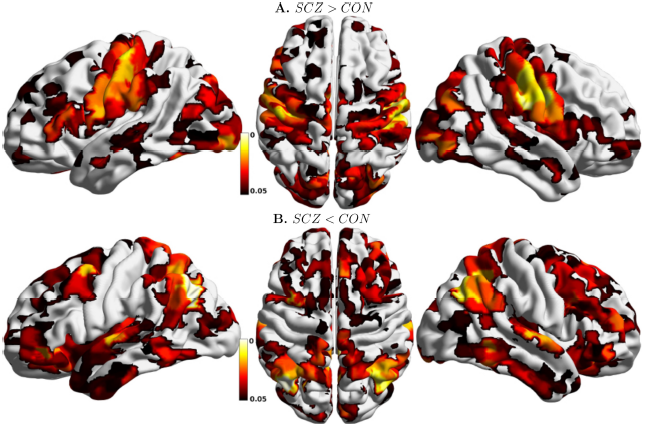
Anatomical distribution of statistically significant node-wise between-group differences in Participation Coefficient, Bonferroni corrected. A. Nodes with higher participation coefficient in SCZ than in CON; B. Nodes with lower participation coefficient in SCZ than in CON.

Significantly larger participation coefficients are observed in sensorimotor, visual and auditory areas of SCZ patients. Lower participation coefficients in patients are observed in frontal and parietal regions, with the most prominent decrease in the temporal primary auditory cortex. Substantial differences are also observed in functional connectivity of areas related with language generation and processing. The anterior part of the Broca area (BA 45), which receives afferent projections from the PFC, shows a decrease in participation coefficient,while the posterior part (BA 44), which has more structural connections with sensory cortices and inferior parietal cortices, shows an increase in PC and appears as a prominent connector hub only in the SCZ group. Connector hubs for the two experimental groups are displayed in the Supplementary Information section (Fig.S2).

## Discussion

One of the main conclusions of this paper is the selective fragmentation of specific functional connectivity modules in schizophrenic patients. This effect was not detected in previous studies, where weaker modularity in SCZ patients was not associated with major disruption or reorganization of the modular structure *per se*. The present study leverages an important methodological development that makes it possible to overcome limitations that may have affected previous investigations. Indeed, it has been recognized that community detection approaches based on the optimization of a global fitness function suffer from a fundamental resolution limit, (Fortunato&Barthelemy 2007). We have recently demonstrated that this limit can prevent detection of important details in the structure and organization of resting state functional connectivity networks, thus hampering detection of differences in community structures of different experimental groups (Nicolini et al. 2016). The finer resolution afforded by Asymptotical Surprise (Nicolini et al. 2017) enables finer grained analysis of resting state brain networks, and provides improved means to assess differences in patient-control studies. Importantly, this new method has been thoroughly validated in synthetic networks endowed with ground-truth modular structures and in functional connectivity networks from healthy subjects (Nicolini et al. 2017), demonstrating superior sensitivity to smaller modules compared to other popular methods, like Newman’s Modularity and InfoMap. Importantly, specificity of Asymptotical Surprise was shown to be in line with or superior to that of resolution-limited methods even in the presence of noise and intersubject variability (Nicolini et al. 2017).

Break-up of functional connectivity modules of SCZ patients was most prominent in primary sensory, auditory and visual areas. Alterations in sensory experience and processing have been documented for a long time (Bleuler 1950; McGhie & Chapman 1961; Chang & Lenzenweger 2005; Dworkin 1994), but studies in schizophrenia have traditionally focused on deficits in higher-order processes such as working memory and executive function. It has been also suggested that bottom-up deficits in cognitive processing may be driven by impairments in basic perceptual processes that localize to primary sensory brain regions (Daniel C. Javitt 2009; Daniel C Javitt 2009). The major reorganization of functional connectivity in sensory areas hereby reported is in keeping with the idea that disorders in schizophrenia may occur already at the level of early sensory processing.

Dysfunction in auditory sensory processing has been consistently observed in schizophrenia with the auditory mismatch negativity (MMN) test (Javitt, Daniel C.; Sweet 2015), and has been related with altered intrinsic and extrinsic connectivity in the Superior Temporal Gyrus (Garrido et al. 2008). Moreover, deficits in sensorimotor gating as measured by the Pre Pulse Inhibition test have been widely documented (Braff & Geyer 1990). Consistent with these observations, our results highlight fragmentation in the organization of the auditory cortex module, resulting in abnormal connectivity within and between early auditory processing areas.

Perceptual deficits have also been documented in the visual system of schizophrenia patients (Butler et al. 2001; Braus et al. 2002). Specifically, alterations have been reported in the magnocellular visual pathway, resulting in deficits in processes such as perceptual closure, object recognition, and reading (Doniger et al. 2002). On the other hand, ERP studies suggest that the ventral stream processing is preserved and that impaired magnocellular dorsal stream schizophrenia may lead to secondary dysregulation of ventral stream object recognition processing (Doniger et al. 2002). Our data provide evidence of the reorganization of functional connectivity between the primary visual cortex and the ventral and dorsal pathways, with stronger connectivity in the ventral stream, leading to merging of primary visual and inferior temporal cortices into a single module, and separation from the dorsal and dorso-parietal visual cortices.

Although most reports focus on the visual and auditory systems, deficits in other sensory systems have also been documented in schizophrenics, including reduced sensitivity to stimulus features (Javitt et al. 1999), impaired 2-point discrimination (Chang & Lenzenweger 2005) and abnormal pain thresholds (Dworkin 1994). This appears consistent with the breakdown of the sensorimotor module reported in the present study.

The effects of abnormal connectivity organization in primary sensory cortices in patients are also apparent in the anatomical distribution of the participation coefficient. This index reflects the balance of within- and between- module connectivity, with higher values denoting regions that project mostly to other modules, and thus play an integrative role within the overall connectivity network. In the healthy brain, high participation is typically found in heteromodal association cortices. By contrast, primary sensory cortices tend to have lower topological centrality (Fornito et al. 2015). In our analyses, we find that sensorimotor and primary visual cortices show a significant increase in participation coefficient in SCZ patients compared to controls. Hence, these regions play an abnormally central role in the network topological integration. Conversely, frontal and parietal cortices, including heteromodal and associative cortices show significantly reduced participation coefficient in patients compared to healthy controls. Interestingly, the primary auditory cortex is also characterized by reduced variety of intermodular connectivity, unlike other primary sensory cortices, and substantial alterations in its connectivity with language processing areas (see below).

It is noteworthy that the overall reduction in connectivity strength observed in patients does not result in unspecific breakdown of functional modules into smaller, more fragmented structures across the whole brain. Indeed, the number of modules in the two groups is comparable, despite the much weaker connectivity strength of the patients’ group. Fragmentation is observed in primary sensory areas, but not in other areas, like the frontal ones. Altogether, we report a reorganization of the connectivity modular structure in SCZ patients, with district specific effects. This is consistent with the idea that the modular structure of a network is determined by the balance between inward and outward links of individual communities, rather than by the average total distribution of edge weights.

Particularly interesting is the reorganization of areas involved in the processing of language. The larger participation coefficient of the Broca area in SCZ subjects causes this area to play the role of connector hub in this population, while parietal heteromodal cortices have a largely reduced integrative role in this population. The Angular Gyrus, a cross-modal hub where converging multisensory information is combined and integrated, plays the role of connector hub in healthy subjects, but not in patients, where it forms a tight community with the Supramarginal Gyrus, a region involved in language perception and processing. Finally, significantly higher values of the participation coefficient are observed in the Heschl gyrus of SCZ patients, an area that has been shown to be overactive during hallucinatory states (Dierks et al. 1999). Hence, we speculate that the abnormal connectivity between language and multisensory integrative areas may be related with the insurgence of auditory hallucinations (“hearing voices”), a hallmark of Schizophrenia.

We note that the groups of subjects included in the study, all selected on the basis of a strict (DSM-IV) Schizophrenia diagnosis, present a wide age distribution at the time of MRI scan. To assess the potential effects of age, we have re-binned the subjects into three subgroups, including subjects up to 25, 35 or over 35 years, respectively. The modular structures for these three subgroups of patients, reported in Fig.S3 in the Supplementary Information, show similar fragmentation and reorganization of sensory cortices, thus indicating that the effects hereby reported are not driven by a restricted age-group of subjects. Moreover, patients in this study are likely to have taken medications, with different histories of pharmacological treatments. Hence, we cannot exclude that differences with respect to the control group may be related with pharmacological treatment, particularly in subjects with long-term exposure to antipsychotic drugs. To mitigate this risk, we have subdivided the patients for whom treatment information is available in different groups based on the difference from time of first treatment and time of MRI study. Specifically, we have taken patients who had the resting state fMRI scan the same year of first treatment (18 subjects), within the year after (35 subjects), or after 5 years. The results are shown in Fig.S4 of the Supplementary Information section. Despite some differences, possibly related with the limited and different number of subjects included in each subgroup, the same general features, including fragmentation of sensory areas, are consistently observed in all subgroups. Altogether, subgroup analysis suggests that the reorganization of functional connectivity hereby reported is not driven by factors like duration of pharmacological treatment. Finally, it has been suggested that increased head motion may affect resting state functional connectivity as measured by functional MRI in certain populations of psychiatric patients, like Autistic Spectrum Disorder subjects. Means and standard deviations of motion parameters extracted with SPM for each subject included in this study are shown in the Supplementary Information section (fig. S5). Analysis of motion parameters did not show any significant difference between groups (z translation: p=0.265; y translation: p=0.525; x translation: p=0.2446; x rotation: p=0.245; y rotation: p=0.652; z rotation: p=0.194), consistent with the fact that patients included in this study were not under acute psychosis. Additionally, it should be noted that fragmentation of modular organization in patients is not widespread throughout the brain, but only observed in specific districts, like the primary sensory cortices, while other modules are almost identical to those of the controls. Hence, it is unlikely that inter-group differences in modular structure be dominated by the effects of differential head motion.

In summary, this study demonstrates previously unreported fragmentation of the modular structure of functional connectivity of primary sensory cortices in SCZ patients. Conversely, prefrontal cortices exhibit normal local organization, despite overall weaker connectivity in SCZ patients. This is interesting, as these areas are thought to be involved in higher cognitive processes, which are profoundly affected by schizophrenia. Our findings support the theory that aberrant connectivity in primary sensory processing may induce deficits that reverberate to higher cognitive functions through a bottom-up process (Daniel C Javitt 2009). Moreover, we report a substantial reorganization of language and speech areas, with an abnormal association of the Supramarginal Gyrus with heteromodal cortices, and an increase in the centrality of the Broca area at the level of network topology.

## Conclusion

In conclusion, we have applied a novel graph theoretical approach, dubbed Asymptotical Surprise, to study the structure of brain functional connectivity networks in a large cohort of Schizophrenia patients. Global and node-wise connectivity parameters showed an overall reduction in connectivity in patients compared to healthy controls, in line with previous studies. The improved resolution afforded by our method revealed substantial reorganization of the modular structure of functional connectivity in patients, with a fragmentation of visual, auditory and sensorimotor cortices. Conversely, prefrontal cortices exhibit normal local organization, despite overall weaker connectivity in SCZ patients. This was perhaps unexpected, as these areas are thought to be involved in higher cognitive processes, which are profoundly affected by schizophrenia. Our findings support the theory that aberrant connectivity in primary sensory processing may induce deficits that reverberate to higher cognitive functions through a bottom-up process. The reorganization of auditory and language modules, and the merger with multimodal association cortices is particularly interesting in the light of the auditory hallucinations often experienced by SCZ patients. Significant changes were observed in the participation coefficient of sensory, visual, and in primary auditory cortices, including the Heschl gyrus, a region critically implicated in auditory hallucinations. This evidence indicates that these regions play a different role in the integration of the network of functional connectivity in the patient’s brain. Previous studies using resolution-limited methods may have failed to detect the abnormal organization of functional connectivity at the scale reported here due to intrinsic methodological limitations. The present approach may provide a novel and powerful tool to study alterations in the brain functional organization in other neuropsychiatric conditions that are thought to be associated with aberrant connectivity.

## Acknowledgments and Disclosures

The Authors wish to thank Dr. Giulia Scuppa and Prof. Enrico Domenici for critically reviewing the manuscript, and Prof. Edward Bullmore and Prof. Nicholas Crossley for providing the brain parcellation template. Data was downloaded from the COllaborative Informatics and Neuroimaging Suite Data Exchange tool (COINS; http://coins.mrn.org/dx) and data collection was performed at the Mind Research Network, and funded by a Center of Biomedical Research Excellence (COBRE) grant 5P20RR021938/P20GM103472 from the NIH to Dr. Vince Calhoun. This project has received funding from the European Union’s Horizon 2020 Research and Innovation Program under grant agreement No 668863.

The authors report no biomedical financial interests or potential conflicts of interest.

